# MIDASim: a fast and simple simulator for realistic microbiome data

**DOI:** 10.1101/2023.03.23.533996

**Authors:** Mengyu He, Ni Zhao, Glen A. Satten

## Abstract

**Background:** Advances in sequencing technology has led to the discovery of associations between the human microbiota and many diseases, conditions, and traits. With the increasing availability of microbiome data, many statistical methods have been developed for studying these associations. The growing number of newly developed methods highlights the need for simple, rapid, and reliable methods to simulate realistic microbiome data, which is essential for validating and evaluating the performance of these methods. However, generating realistic microbiome data is challenging due to the complex nature of microbiome data, which feature correlation between taxa, sparsity, overdispersion, and compositionality. Current methods for simulating microbiome data are deficient in their ability to capture these important features of microbiome data, or can require exorbitant computational time.

**Methods:** We develop MIDASim (<u>MI</u>crobiome <u>DA</u>ta <u>Sim</u>ulator), a fast and simple approach for simulating realistic microbiome data that reproduces the distributional and correlation structure of a template microbiome dataset. MIDASim is a two-step approach. The first step generates correlated binary indicators that represent the presence-absence status of all taxa, and the second step generates relative abundances and counts for the taxa that are considered to be present in step 1, utilizing a Gaussian copula to account for the taxon-taxon correlations. In the second step, MIDASim can operate in both a nonparametric and parametric mode. In the nonparametric mode, the Gaussian copula uses the empirical distribution of relative abundances for the marginal distributions. In the parametric mode, an inverse generalized gamma distribution is used in place of the empirical distribution.

**Results:** We demonstrate improved performance of MIDASim relative to other existing methods using gut and vaginal data. MIDASim showed superior performance by PERMANOVA and in terms of alpha diversity and beta dispersion in either parametric or nonparametric mode. We also show how MIDASim in parametric mode can be used to assess the performance of methods for finding differentially abundant taxa in a compositional model.

**Conclusions:** MIDASim is easy to implement, flexible and suitable for most microbiome data simulation situations. MIDASim has three major advantages. First, MIDASim performs better in reproducing the distributional features of real data compared to other methods at both presence-absence level and relative-abundance level. MIDASim-simulated data are more similar to the template data than competing methods, as quantified using a variety of measures. Second, MIDASim makes few distributional assumptions for the relative abundances, and thus can easily accommodate complex distributional features in real data. Third, MIDASim is computationally efficient and can be used to simulate large microbiome datasets.

## 1 Introduction

The human microbiota and its associated microbiome play a fundamental role in many diseases and conditions, including obesity [1], inflammatory bowel disease (IBD) [2], preterm birth [3], autism [4] and cancers [5, 6]. Advances in sequencing technologies, especially 16S rRNA sequencing, now allow rapid and simultaneous measurement of the relative abundance of all taxa in a community. This has led to a growing number of epidemiological and clinical studies to measure the association between the microbiome and traits of interest, sometimes with complex study designs and research questions.

Although microbiome data is increasingly available, statistical analysis remains challenging. Microbiome data have special characteristics that are difficult to model analytically, including sparsity (the majority of taxa are not present in a sample), overdispersion (the variance of read counts is larger than what is assumed from the usual parametric models), and compositionality (the read counts in a sample sum to a constant). There is little consensus among researchers on how microbiome data should be analyzed, and new methods are being regularly developed, both for identifying individual taxa that associate with diseases [7, 8, 9, 10, 11, 12, 13], and for understanding the community-level characteristics that relate to clinical conditions [14, 15, 16].

Simulating realistic microbiome data is essential for the development of novel methods. To establish the validity of a new method and prove it outperforms existing ones, researchers rely on simulated data in which the true microbiome/trait associations are known. Ideally, the simulated data should be similar to real microbiome data for the simulation studies to be trustworthy. How-ever, simulating realistic microbiome data is made difficult by the same challenges as analyzing microbiome data: sparsity, overdispersion and compositionality. Further, the distribution of counts for each taxon are highly skewed and correlated in a complex way. For these reasons, most simulation methods are based on using a *template* microbiome dataset and generate simulated data that is “similar” to the template data in some way.

Several approaches have been proposed for simulating microbiome data. Among them, some methods impose strong parametric assumptions so that the simulated microbiome data share similar dispersion of real data. For example, the Dirichlet-Multinomial (D-M) distribution, in which the taxa counts are generated from a multinomial distribution with proportion parameters provided by a Dirichlet prior [17], is frequently used in simulating microbiome data. The hyper-parameters of this D-M model are often estimated from real data so that the simulated data share similar dispersion. Another method, MetaSPARSim [18], uses a gamma-multivariate hypergeometric (gamma-MHG) model, in which the gamma distribution models the biological variability of taxa counts, accounting for overdispersion, and the MHG distribution models technical variability originating from the sequencing process.

Although the D-M model and the MetaSPARSim model address the compositional feature by either the multinomial or the hypergeometric distribution, they do not attempt to match the correlation structures in the simulated data with those found in the real data. One recently developed approach that does attempt to model between-taxa correlations is SparseDOSSA (Sparse Data Observations for the Simulation of Synthetic Abundances) [19]. This hierarchical model makes assumptions about both the marginal and joint distributions of the relative abundances of a set of taxa. For the marginal distribution, SparseDOSSA assumes a zero-inflated log-normal model for the relative abundance of each taxon and then imposes the compositional constraint. Parameters in the zero-inflated log-normal marginal are estimated through a penalized Expectation-Maximization (EM) algorithm from a template dataset. Unfortunately, the penalized EM algorithm for estimating hyper-parameters is computationally expensive, especially when a large number of taxa exist in the data. For example, fitting SparseDOSSA model to a modest-sized dataset with sample size of 79 and number of taxa = 109 takes more than a day (≈ 27.8 hours) on a single Intel “Cascade Lake” core [19]. To partially compensate for this drawback, SparseDOSSA provides fitted models that were previously trained by the developers and that users can use directly, which is only useful if the developer-provided fits resemble the data users wish to generate. Moreover, SparseDOSSA removes rare taxa that appear in fewer than 4 samples by default, thus failing to accommodate the possibility that rare taxa are of interest in the simulation studies.

Recently, deep neural networks have also been used in simulating microbiome data, notable examples being MB-GAN [20] and DeepMicroGen [21]. MB-GAN employs a deep generative adversarial network (GAN) to autonomously learn from actual microbial abundances, obviating the need for explicit statistical modeling assumptions. DeepMicroGen, tailored for longitudinal microbiome studies, utilizes a bidirectional recurrent neural network (RNN)-based GAN to impute missing data by exploiting temporal relationships between samples. Although these deep neural network models show promise over conventional statistical models in capturing microbiome data’s complex structure, their practical application is challenging. Issues include the difficulty in tailoring simulations to specific alterations in data structure (e.g., changes in relative abundances), and severe computational issues (see https://github.com/zhanxw/MB-GAN/blob/master/code_check_convergence/plot_logs_convergence_check.ipynb). Consequently, these methods were not included in our comparative analyses.

Considering the drawbacks of existing approaches, a method that can flexibly capture the distributional and correlation structure of microbiome data would greatly benefit the research community. Here, we develop a fast and simple <u>MI</u>crobiome <u>DA</u>ta <u>Sim</u>ulator (MIDASim) for generating realistic microbiome data that capture the correlation structure of taxa of a template microbiome dataset in both the presence-absence structure and the relative abundances. MIDASim can operate in two modes: parametric and nonparametric. In nonparametric mode, all quantities are calculated using their empirical distributions in the original data. In parametric mode, we use an inverse generalized gamma distribution to model the relative abundances; this model is fit using a novel method-of-moments approach. We show that the resulting distribution gives good agreement with the datasets we analyze here, for both low and high prevalence taxa. The parametric mode is primarily designed for simulation studies where we want to make changes to the log-mean relative abundance so that we can assess the performance of methods that look for differentially abundant taxa in log-linear models such as the compositional model. Using simulations, we show that MIDASim in either mode generates data that are more similar to the template data, as measured by multiple metrics, than competing methods. MIDASim is implemented as an R package (https://github.com/mengyu-he/MIDASim).

## 2 Results

### 2.1 The MIDASim approach

MIDASim simulates microbiome data using a two-step approach. The first step generates the presence-absence status for taxa in each sample by simulating correlated binary data from a probit model with a correlation structure chosen to match the empirical correlation in the template data. The second step generates relative abundance and count data for non-zero taxa from a Gaussian copula model.

This model allows for separate fitting of each taxon’s relative abundance marginal distribution and the inter-taxa correlations. For taxon-taxon correlation, MIDASim employs a rank-based approach to accurately mirror the empirical correlations observed in the template data, effectively managing zero counts. Regarding the marginal distribution, MIDASim offers two options: using the taxon-specific empirical distribution (nonparametric mode) or sampling taxon relative abundances from an inverse generalized gamma distribution (parametric mode). This flexibility enables MIDASim to capture the complex distributional characteristics often present in real data.

MIDASim also allows the user to change the library sizes, taxon relative abundances or the proportion of non-zero cells, and these features may depend on covariates such as case/control status. MIDASim is computationally efficient and can be used to simulate large microbiome datasets in a fast and simple fashion.

### 2.2 Simulation setup

We compared MIDASim in both parametric and nonparametric mode to three competing methods (the D-M method, MetaSPARSim and SparseDOSSA) and evaluate how well the simulated data reporduce the characteristics of the template data. We use two datasets from the Integrative Human Microbiome Project (HMP2) [22] as the template data: a vaginal microbiome dataset from Multi-Omic Microbiome Study: Pregnancy Initiative (MOMS-PI) project, and a gut microbiome dataset from the Inflammatory Bowel Disease Multi-omics Database (IBDMDB) project [23]. These two datasets represent microbiomes from two body sites that are frequently studied in the literature. They are also distinct in their characteristics, and thus provide a comprehensive assessment of the proposed method. For example, the vaginal data is notably sparse, comprised of 95.25% zeros. In contrast, the gut data is less sparse, comprised of 85.09% zeros. Both datasets feature taxa that are OTUs; the IBD data are classified at the genus level, while the MOMS-PI data are classified to the species level using a “best guess” approach. Moreover, the coefficient of variation (CV) of vaginal data is 40.77, while that of the gut data is 10.76, indicating that the vaginal data is more over-dispersed. We compared the four methods using two aspects of performance: how well the simulated data matched the template data, and the computational effort required to fit and generate a simulated dataset. Further details on the statistical procedures used can be found in Supplemental text (Section: Statistical Analyses).

Before fitting MIDASim, we lightly filtered the two template datasets. For quality control, we removed samples with library size *<* 3000. To allow comparison with SparseDOSSA, we removed taxa that were present in fewer than 4 samples, a requirement of SparseDOSSA. MOMS-PI is a longitudinal study with repeated vaginal samples; we kept only first-visit samples to avoid repeated measures. The only filtering used for the IBD data was that required by SparseDOSSA. After filtering, 517 samples and 1146 taxa were preserved in the vaginal MOMS-PI dataset; the gut IBD dataset comprised 146 samples and 614 taxa. This filtering also slightly decreased the zero proportions in the template datasets. Specifically, in the IBD dataset, the zero proportion was reduced from 89.69% to 85.09% following the filtering. Similarly, for the MOMS-PI dataset, the zero proportion decreased from 96.97% to 95.25%. We ignored covariates such as gender or location of biopsy collection to focus only on reproducing the microbiome datasets as closely as possible, the goal of all methods considered here. In our simulations, the library sizes for datasets generated using the D-M method and MetaSPARSim were the same as those in the original data. For SparseDOSSA, the library sizes were generated from a log-normal distribution parameterized by mean and standard deviation of log counts in the original data, as recommended in their original publication. To facilitate a comparison of the methods, all simulated counts were transformed to relative abundances.

### 2.3 MIDASim outperforms existing methods in reproducing distributional features of microbiome data

The PCoA plots in Figure 1 provide a simple visualization of the similarities between the original data and the simulated data by MIDASim (in both nonparametric and parametric modes), the D-M method, MetaSPARSim, and SparseDOSSA for the IBD data and MOMS-PI data. For both datasets, after ordination, the data simulated from MIDASim looked similar to the template data, using either the (presence-absence-based) Jaccard distance (Figure 1 A,C) for nonparametric, (E,G) for parametric or (relative abundance-based) Bray-Curtis distance (Figure 1 B,D) for nonparametric, (F,H) for parametric. Conversely, for both data templates, data simulated by the D-M method, MetaSPARSim, SparseDOSSA all appear to be underdispersed in the first two principal coordinates (Figure 1 I,K,M,O,Q,S) using the Jaccard distance. For the IBD data, data simulated using D-M and MetaSPARSim appeared easily distinguishable from the original data when the Bray-Curtis distance was used (Figure 1 J, N). For both the IBD and the MOMS-PI data, we also see clear underdispersion in data simulated using D-M (Figure 1 J,L). To allow visual comparison between the template data and multiple datasets simulated by MIDASim, in Figure S1 we also give a probability density map of data generated using MIDASim, constructed using 20 simulated datasets. In general, the agreement between the observed and expected values is good.

**Figure 1:**
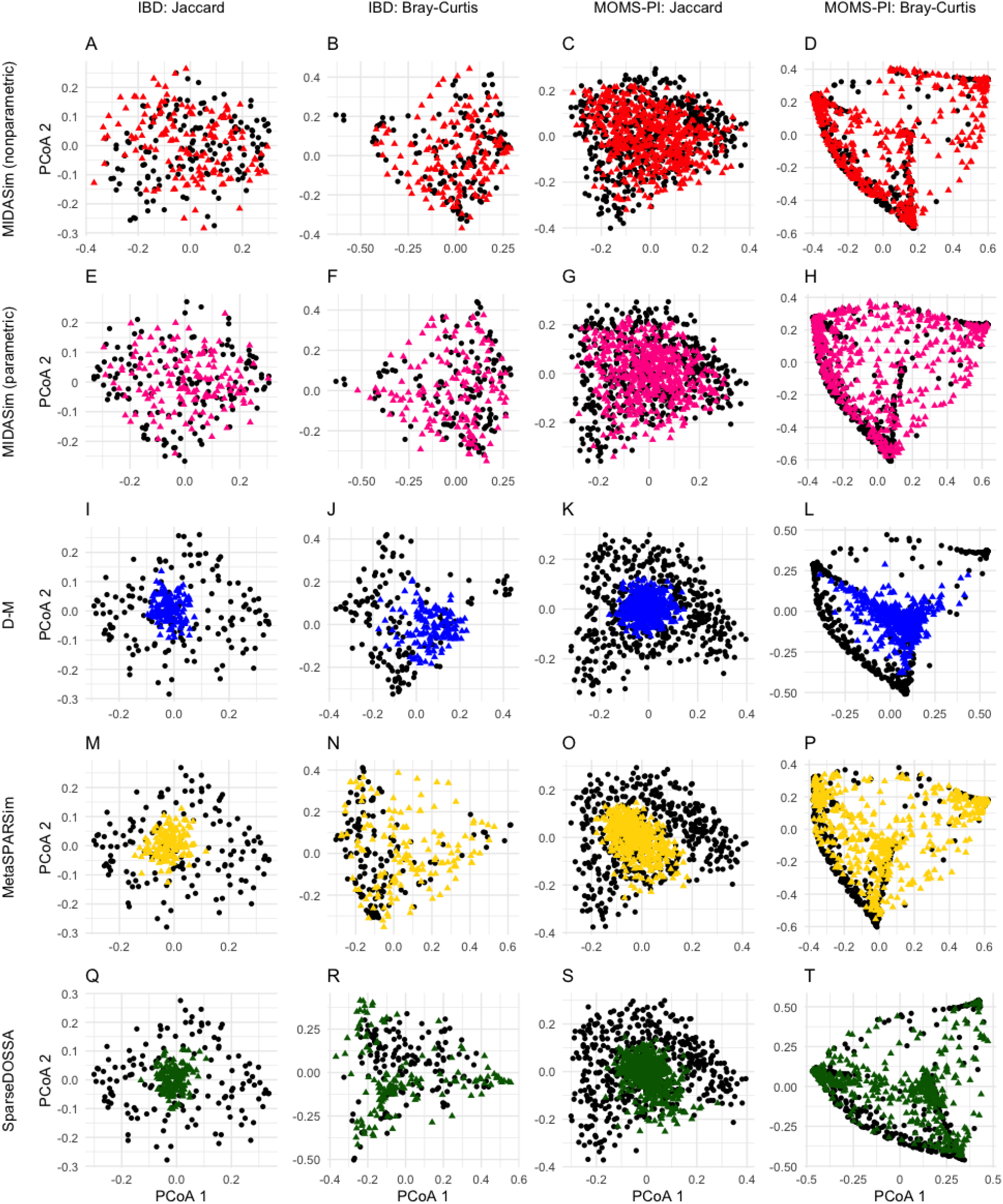
Principal Coordinates plots (PCoA) of the simulated and original community. Each row corresponds to one method. The left two columns are the plots for the IBD data, and the right two columns are the plots for the MOMS-PI data. Black points: samples from original data. Colored points: samples from the simulated data with red being MIDASim with nonparametric model, yellow being MIDASim with parametric model, blue being D-M, pink being MetaSPARSim, and green being SparseDOSSA.

The visual impressions of beta diversity in figures Figure 1 and Figure S1 are confirmed in Table 1, where we test whether the template and simulated data are significantly different in beta diversity using PERMANOVA [24]. For tests using the Jaccard distance, the *p*-values for MIDASim in nonparametric mode were consistently high (indicating no detected difference between simulated and template data); in parametric mode, MIDASim had a significant difference for the MOMS-PI data but not the IBD data. For all other methods PERMANOVA found highly significant differences between the simulated and template data with the single exception of SparseDOSSA applied to the IBD data using the Jaccard distance. Note that when using the Bray-Curtis distance, only MIDASim in nonparametric mode could produce data that was not easily differentiated from the template data by PERMANOVA.

**Table 1:** Average *p*-value (from 20 replicates) for tests comparing alpha and beta diversities of simulated data and template data. The significance level is 0.05.

| Data | Method | Beta-Diversity* |  | Alpha-Diversity** |  |  |  |
| --- | --- | --- | --- | --- | --- | --- | --- |
|  |  | Jaccard | Bray-Curtis | Richness t | Richness KS | Shannon t | Shannon KS |
| IBD | MIDASim | 0.9993 | 1.0000 | 0.6644 | 0.2557 | 0.6047 | 0.6627 |
|  | MIDASim (parametric) | 0.5856 | 0.8118 | 0.4916 | 0.1960 | 0.3306 | 0.2565 |
|  | D-M | 0.0090 | < 0.0001 | 0.3303 | < 0.0001 | < 0.0001 | < 0.0001 |
|  | MetaSPARSim | 0.0340 | < 0.0001 | 0.3102 | < 0.0001 | 0.0078 | < 0.0001 |
|  | SparseDOSSA | 0.7972 | < 0.0001 | 0.0569 | < 0.0001 | < 0.0001 | < 0.0001 |
| MOMS-PI | MIDASim | 0.5793 | 0.8617 | 0.6252 | 0.0019 | < 0.0001 | < 0.0001 |
|  | MIDASim (parametric) | 0.0058 | 0.0010 | 0.0495 | 0.1607 | < 0.0001 | < 0.0001 |
|  | D-M | < 0.0001 | < 0.0001 | 0.0028 | < 0.0001 | < 0.0001 | < 0.0001 |
|  | MetaSPARSim | < 0.0001 | < 0.0001 | 0.6341 | < 0.0001 | < 0.0001 | < 0.0001 |
|  | SparseDOSSA | < 0.0001 | < 0.0001 | < 0.0001 | < 0.0001 | 0.0002 | 0.0015 |

\* Beta-diversity comparisons were conducted using PERMANOVA.
\*\* Alpha-diversity comparisons were conducted using both $t$ -test and the Kolmogorov-Smirnov (KS) test.

To compare the performance of all methods in terms of beta dispersion, in Figure 2 we compare the empirical cumulative distribution function (CDF) of the distance between each sample and the group centroid in the simulated data to this CDF in the template data. These distances were calculated using the `betadisper` function in the R package `vegan`. If the simulated data are similar to the template data, the CDF of distances-to-centroids in the simulated data should resemble that of the template data. These CDFs are shown in Figure 2 for Jaccard and Bray-Curtis distances, for the IBD and MOMS-PI data. The CDFs datasets simulated by the D-M method, MetaSPARSim, and SparseDOSSA are noticeably dissimilar to the CDFs of the template data; this dissimilarity is confirmed by extremely small Kolmogorov-Smirnov two-sample test *p*-values reported in the figure. The range of distances to centroids in the data simulated by the D-M method and Sparse-DOSSA is smaller compared to the real data in every scenario, indicating a smaller dispersion overall. For the IBD data, the MIDASim-simulated data (both modes) follow the template data closely in dispersion in both Jaccard and Bray-Curtis distances. For the MOMS-PI dataset, the non-parametric MiDASim generated data exhibiting a dispersion profile similar to the template data when evaluated using the Jaccard distance, but not the Bray-Curtis distance. Conversely, the parametric MiDASim yielded data with significant differences in both Jaccard and Bray-Curtis distance measures. However, panel C and D of Figure 2 show the MIDASim results (especially in nonparametric mode) are clearly closer to those of the template data than the other methods are.

**Figure 2:**
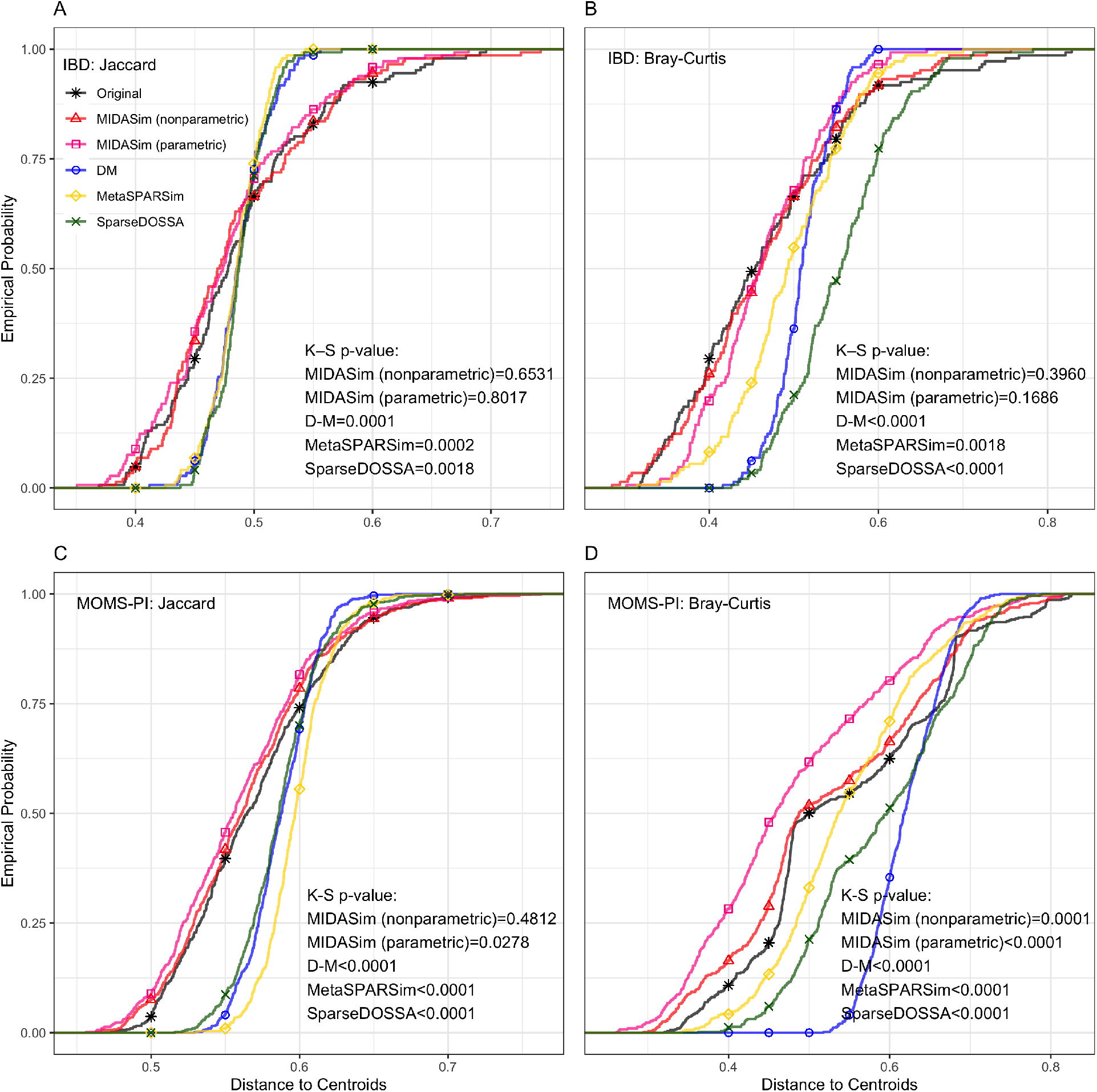
Empirical cumulative distribution function of distances to centroids

Figures S2 and S3 display the results of t-distributed Stochastic Neighbor Embedding (t-SNE) and Uniform Manifold Approximation and Projection (UMAP) analyses, applied to simulated and template data using Jaccard and Bray-Curtis distances using multiple methods. These visualizations corroborate the findings from the PCoA plot, demonstrating that data generated by MIDASim more closely resemble the template data compared to those from alternative methods.

Table 1 and Figure 3 present comparisons of two alpha diversity measures: species richness and the Shannon index. We employed the Welch *t*-test to compare the mean alpha diversities and the Kolmogorov-Smirnov two-sample test for differences in their distributions. Table 1 reports the average *p*-values obtained from 20 simulated datasets for each method. In the IBD data analysis, all methods successfully reproduced mean richness (indicated by Welch *t*-test *p*-values *>* 0.05). For the MOMS-PI data, only MIDASim (in nonparametric mode) and MetaSPARSim produced mean richness values not significantly different from the template data. A different perspective emerges when analyzing the entire distribution of sample richness using the Kolmogorov-Smirnov test. Here, only MIDASim (in both modes) generated data with richness distribution indistinguishable from the IBD data, and only MIDASim in parametric mode achieved this for the MOMS-PI data. Regarding the Shannon index, MIDASim (in both modes) was the only method to successfully generate data resembling the template IBD data in both mean and distribution. However, for the MOMS-PI data, no method could replicate the Shannon index of the template data. It is noteworthy that, even when MIDASim indicated significant differences sometimes, its *p*-values were often larger than those of competing methods. Figure 3 also illustrates the alpha diversities for a single dataset from each simulation method, where MIDASim more closely matches the template data’s alpha diversity. Additionally, the alpha diversity of MIDASim in parametric mode is typically less variable than in nonparametric mode, potentially explaining its relative performance in beta diversity.

**Figure 3:**
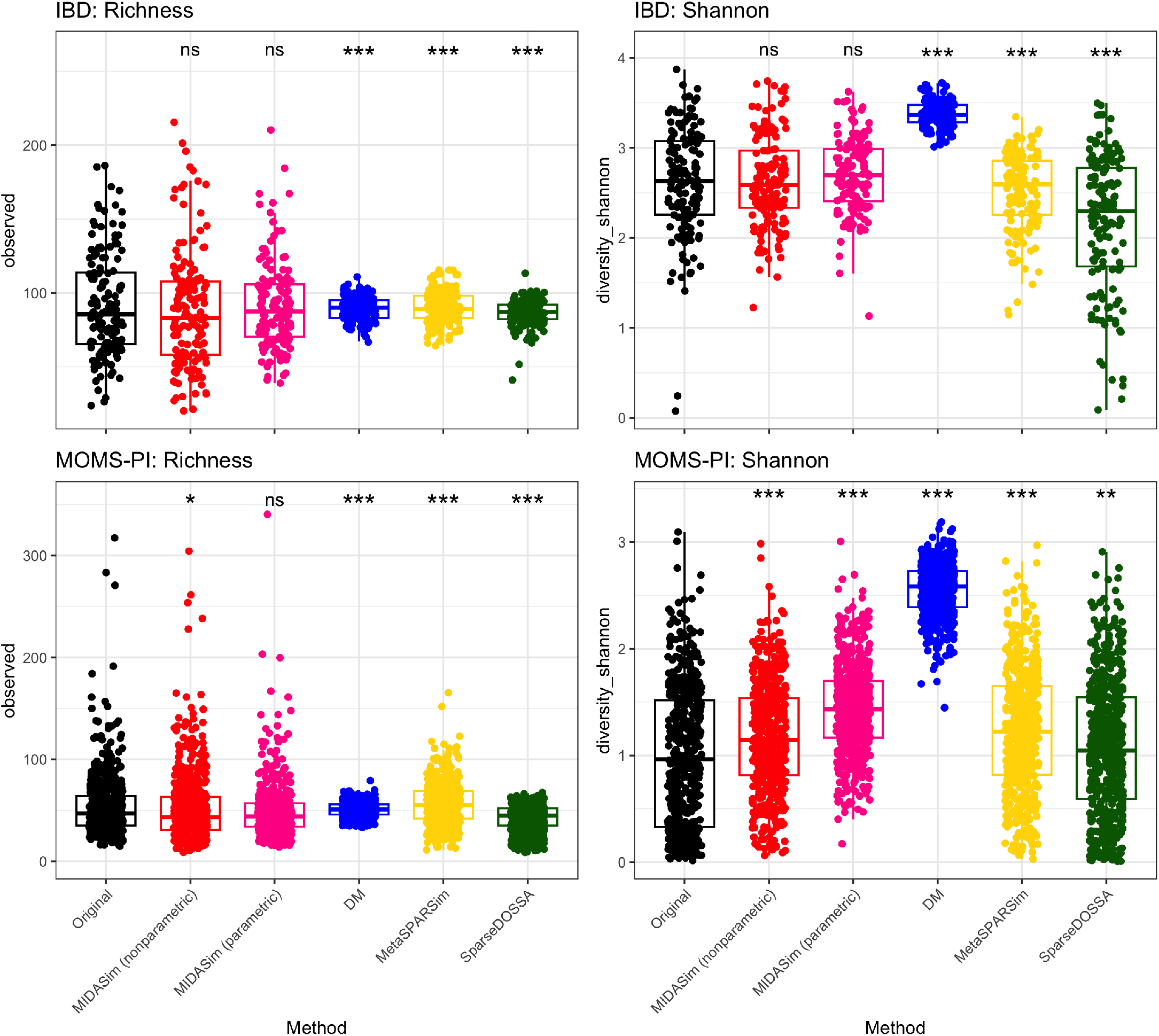
Alpha diversities (Richness and Shannon Index) of original and a single simulated dataset for each of four simulation methods. Asterisks indicate significance levels of KS-test p-values comparing the simulated data with that in the template data, as shown in Table 1: ns (*p >* 0.05), * (*p <* 0.05), ** (*p <* 0.01), *** (*p <* 0.0001).

We also applied MIDASim to the un-filtered datasets to assess its performance when very rare taxa are present. Including all taxa, the IBD data comprised 908 taxa for 146 subjects, and the MOMS-PI data comprised 1839 taxa for 517 subjects. We compared the alpha and beta diversities between the template data and the MIDASim simulated data in Table S1. The result remains consistent with scenarios where extremely rare taxa are excluded.

### 2.4 MIDASim can be used for assessing newly designed statistical tools

To demonstrate the capability of MIDASim for evaluating newly developed statistical tools, we used MIDASim to generate realistic microbiome data that included taxa with relative abundances that varied with categorical covariates. We used the IBD data [23] as the template, resulting in the simulation of 614 taxa across *n* independent samples. A more detailed description of the simulation can be found in Section 4.5. Briefly, we generated a dichotomized covariate *X*_1_ that affected the relative abundance of either 10 or 20 “causal” taxa, randomly selected among the 100 taxa having the highest relative abundances. We generated a second covariate *X*_2_ that affected a second group of 10 taxa selected in the same way, such that there were always 5 taxa affected by both covariates. We assumed *X*_2_ had a fixed effect on relative abundances, but varied the effect of *X*_1_ according to a parameter that measures the effect size. The precise effect of the covariates is given in Equations (9) and (10). *X*_1_ and *X*_2_ are simulated to be balanced. Note that although only a subset of taxa are directly affected by our covariates, the relative abundances of all other taxa are modified due to the compositional constraint that relative abundances sum to one.

We used data simulated with MIDASim to evaluate seven existing methods that can measure the association between *X*_1_ and each taxon while adjusting for *X*_2_. These methods are: (1) Analysis of Compositions of Microbiomes with Bias Correction (ANCOM-BC) [25], (2) an updated version of ANCOM-BC which additionally accounts for taxon-specific bias (ANCOM-BC2) [26], (3) the original Linear Decomposition Model (LDM) as proposed in [11], (4) an updated LDM version incorporating the centered log-ratio transformation [27], (5) the Linear models for Differential Abundance analysis (LinDA) [28], (6) the Logistic Compositional Model (LOCOM) [13], and (7) the Zero-Inflated Quantile approach (ZINQ) [29]. Notably, ZINQ and the original LDM is designed to test differences in relative abundances, while the other methods are tailored for the compositional null hypothesis. Our analysis was restricted to taxa present in at least 20% of the samples.

Figure 4 presents the False Discovery Rate (FDR) at a nominal 0.2 rate for all evaluated methods when *n* = 200. Results for *n* = 100 are analogous and have been omitted for brevity. Unsurprisingly, ZINQ and the original LDM model exhibit a notably inflated FDR, as they test the hypothesis of any difference in relative abundance. In MIDASim-simulated data, changes in the abundance of one taxon can influence the relative abundances of others due to compositional constraints, as described in Equations (9) and (10). Among the remaining methods, which were designed to test the compositional hypothesis, LOCOM shows the best FDR control, followed by LDM-CLR, LinDA and the original ANCOM-BC. To our surprise, the ANCOM-BC2 reports worse FDR control compared to the original ANCOM-BC, possibly due to the difficulty in addressing the taxon-specific bias factor. These findings underscore the efficacy of MIDASim in generating datasets conducive to the evaluation of novel statistical models.

**Figure 4:**
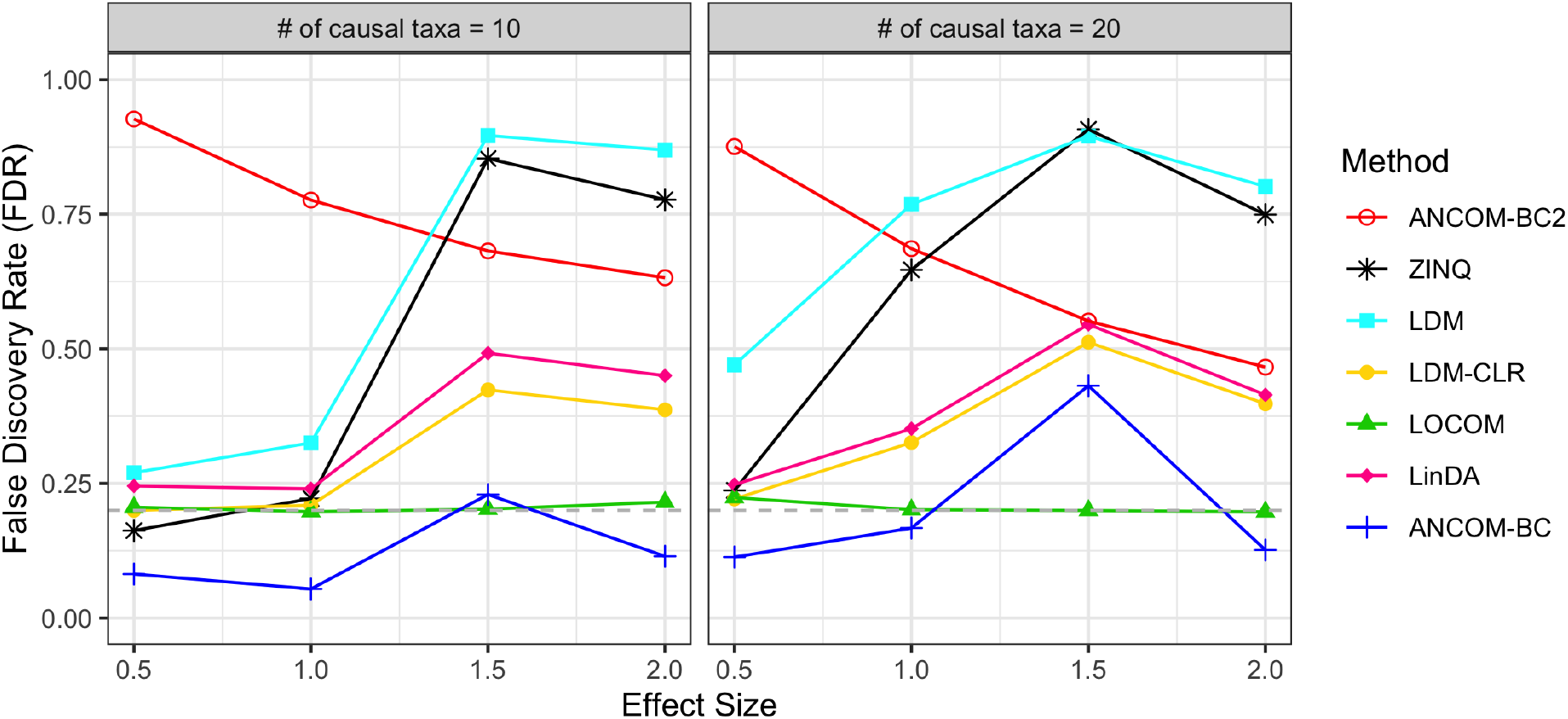
False discovery rate assessment of seven differential abundance analysis methods using MIDASim simulated datasets. Sample size *n* = 200. Effect size is the value of *β*_1_ in Equation 9 and Equation 10. Grey dashed line: FDR = 0.2 reference line.

### 2.5 MIDASim is computationally efficient

We compared the computational time that each method takes to fit its proposed model to the template IBD and MOMS-PI datasets and to simulate one dataset of the same size, which was summarized in Table 2. The computational time was evaluated on an Intel Quad core 2.7GHz processor, with 8GB memory. Comparing the total time used, MIDASim is one of the fastest, especially for the large MOMS-PI dataset. For model fitting, MetaSPARSim is the fastest, but it is very slow in generating new data. For simulating new data after fitting, D-M is the fastest. The computation time of SparseDOSSA for fitting the model depends on the number of iterations in its EM algorithm. We found it took more than 3 hours to fit SparseDOSSA to either the IBD or MOMSPI dataset, making it hard to use in practice; the pre-trained models can be used if faster results are needed, but then a user-selected template dataset cannot be used. Discounting the time required for model fitting, MIDASim, D-M and SparseDOSSA all can generate replicate datasets quickly; MetaSPARSim is the only outlier in this regard.

**Table 2:** Computation time (seconds) required to fit the template data, and to simulate a new dataset with the same library size. Simulating time is the average time over 20 replicates of generating datasets of the same size as the real data. Total time is the sum of fitting and simulating times.

| Method | IBD |  |  | MOMS-PI |  |  |
| --- | --- | --- | --- | --- | --- | --- |
|  | Fitting | Simulating | Total | Fitting | Simulating | Total |
| MIDASim (non-parametric) | 25.5 | 2.5 | 28.0 | 162.0 | 15.3 | 177.6 |
| MIDASim (parametric) | 19.4 | 1.8 | 21.2 | 306.8 | 16.9 | 323.7 |
| D-M | 25.0 | 0.3 | 25.3 | 308.4 | 2.2 | 310.6 |
| MetaSPARSim | 7.4 | 144.9 | 152.3 | 41.3 | 469.4 | 510.7 |
| SparseDOSSA | 10812.6 | 0.8 | 10813.4 | 11792.5 | 5.2 | 11797.7 |

## 3 Discussion

Simulating realistic microbiome datasets is essential for methodology development in microbiome studies. However, this task is surprisingly difficult due to the complexity of microbial relative abundance data. Existing parametric microbiome data simulators facilitate easy simulation of microbiome data in a controlled manner. However, they often fall short in generating realistic correlation structures and accurately reproducing the marginal distributions. In contrast, deeplearning-based methods show promise in effectively modeling complex correlation structures and generating appropriate marginal distributions of microbiome data. However, they typically encounter practical application challenges and are often not user-friendly for generating microbiome data with controlled variations. Here we adopt an empirical approach, using the presence-absence correlation structure of the original data (through a smoothed tetrachoric correlation matrix) and the empirical correlation matrix of relative abundances (using a Gaussian copula model). The use of a Gaussian copula model allows us to closely match the marginal distribution of taxon-specific relative abundances found in the template data, either by using the empirical distribution or by fitting an inverse generalized gamma distribution. Although these assumptions are not based on any underlying model of what microbiome data ‘should’ look like, this approach is fast, easily implemented and appears to reproduce data from a template microbiome dataset better than the existing methods we considered here.

MIDASim can operate in two modes: parametric or nonparametric. Our simulations show that data generated using the nonparametric mode is closer to the template data than data generated using the parametric mode. Thus, if the only goal is to reproduce template data, nonparametric mode should be used. However, data generated in parametric mode may be more useful for simulation studies, since the parametric model correctly adjusts other parameters such as the proportion of non-zero cells when a user changes the taxon mean relative abundances or library sizes. Since it can be difficult to correctly adjust these parameters in nonparametric mode, we strongly suggest using parametric mode for simulations of the type we illustrate in section 2.4. Further, our simulations show that even though data generated in nonparametric mode is more faithful to the template data, the data generated in parametric mode is generally more faithful to the original data than the other methods we studied here.

Although MIDASim does not explicitly support modeling covariates that affect mean relative abundance, it is fairly easy to handle discrete covariates such as case/control status or multiple arms of the same experiment by (1) generating correlations for zero-one and quantitative data from the template data, and then (2) using these correlations to generate data for each covariate group using, say, a different vector of mean relative abundances. We showed here that simulation studies of existing methods using this approach have appropriate false-discovery rate (FDR) when MIDASim-generated data is used.

Compared to competing methods, MIDASim offers users greater flexibility in changing parameters than the Dirichlet-Multinomial model and MetaSPARSim, while providing a better fit to data even in its parametric mode. Further, MIDASim runs much faster than computationally intensive approaches such as sparseDOSSA and the deep-learning-based approaches. The main disadvantages of MIDASim come primarily from its empirical approach; it makes no attempt to base simulations on knowledge of microbiology or microbial ecology, but instead attempts to empirically model observed patterns of correlation. There are several areas where MIDASim could be improved. For example, in its current version, it cannot leverage the correlations found in longitudinal data as DeepMicroGen can. Second, it assumes that the observed correlations are not functions of extra covariates. The use of underlying Gaussian models for generating both presence/absence and qualitative data imposes some limitations on the possible correlation structures available in MIDASim. This last objection could be partially ameliorated for the presence/absence data by providing alternative models to the approach in Equations (1) and (2). The user could then choose the model that best agreed with the template data. Similarly, it may be possible to find a better model for relative abundance data than the generalized gamma, and future revisions could include different choices for this distribution. Additionally, the parametric mode is set up to test the compositional null hypothesis; future revisions could include parametric models that are appropriate for other hypotheses. Finally, we hope to extend MIDASim to handle continuous covariates in a future revision.

## 4 Materials and methods

We assume a template dataset having *n* samples and *J* taxa such that each taxon is present in at least one sample. For sample *i* and taxon *j*, let *C*_*i j*_ denote the observed count, 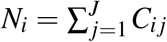 denote the observed library size, *π*_*i j*_ denote the observed relative abundance (*π*_*ij*_ = *C*_*i j*_*/N*_*i*_), and let presence-absence indicator *Z*_*i j*_ = 𝕀(*C*_*i j*_ > 0) where 𝕀(*S*) = 1 if *S* is true and 0 otherwise. We and let ***p*** and ***δ*** be the *J*-dimensional vectors having elements 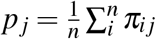 and 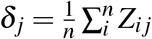 respectively.

We let **C, Z** and **π** represent the *n × J* matrices of the read counts, presence-absence and the relative abundances of all taxa in the template data, respectively. Corresponding quantities for the simulated data are denoted by a tilde, e.g. 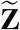 is the presence-absence indicator in the simulated data. We also use a ‘dot’ notation to refer to the *i*^th^ row or *j*^th^ column of matrix *M* as *M*_*i*·_ or *M*_·*j*_, respectively.

MIDASim is a two-step procedure for generating count and relative abundance data. The first step generates binary presence-absence indicators having correlation structure similar to the template presence-absence data ***Z***. This step determines which cells have zero counts in the simulated data. The second step is to fill the non-zero cells from step 1 using a Gaussian copula model fitted to the observed values **π**. In this step, MIDASim provides two options for modeling the marginal distribution of each taxon: a nonparametric mode that uses the empirical distribution, and a parametric mode employing a three-parameter generalized gamma distribution. These modes are accordingly designated as “non-parametric” and “parametric” approaches, based on the marginal distribution choice in this step. We next describe each step in detail for the nonparametric mode; in Section 4.3 we describe the differences when the parametric mode is used.

### 4.1 Step 1: generate presence-absence data

The goal of step 1 is to generate presence-absence data 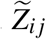 having correlation and marginal means that match the presence-absence structure in the target data. MIDASim uses a threshold model with underlying multivariate normal data *D*_*ij*_ having mean *θ*_*j*_ + *η*_*i*_ and variance-covariance matrix ***ρ*** in such a way that *Z*_*ij*_ = 1 corresponds to *D*_*ij*_ ≥ 0. To accomplish this, we choose *θ*_*j*_ and *η*_*i*_ to jointly solve

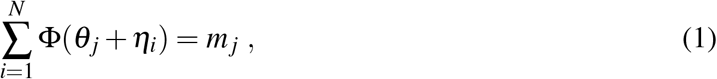

and

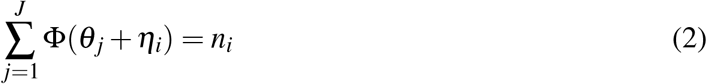

where 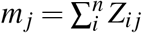 is the number of non-zero cells in the data from the *j*^th^ taxon, 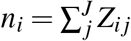 is the number of non-zero cells for the *i*^th^ observation, and Φ(·) and Φ^−1^(·) are the CDF and quantile function of the standard normal distribution respectively. These equations are iterated alternately, starting from the initial values *η*_*i*_ = 0 and *θ*_*j*_ = Φ^−1^(*Z*_· *j*_).

To estimate *ρ*, we first calculate the tetrachoric correlation matrix, denoted by ***ζ***, using the approach of [30]. We smooth ***ζ*** to be positive definite using the function `cor.smooth()` in R package `psych` [31], and denote the resulting correlation matrix 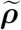. We then sample values 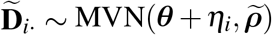 and take 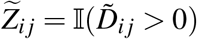.

### 4.2 Step 2: generate relative abundance and count data

We generate relative abundance data using a Gaussian copula model, which allows us to incorporate dependence between taxa while specifing a marginal distribution for each taxon that matches the observed distribution of non-zero relative abundances for that taxon.

In order to allow for the possible generation of non-zero relative abundances for taxa that are observed to have zero counts, we must include the zero cells when we specify the correlation structure of the Gaussian copula. To accomplish this, we use a rank-based approach based on the relationship between the Pearson and Spearman correlations for normally distributed data [32]. This approach does not require us to know the values we would have obtained for an empty cell, had that cell not been empty; our only assumption is that the relative abundances of the zero cells are smaller than those of the cells having non-zero counts. In particular, to specify the correlation of the underlying Gaussian model, we calculate Spearman’s rank correlation ϕ for the observed relative abundance values. When calculating the rank correlation, we consider the zero cells to be tied, and then break these (and any other) ties by a random ordering. For the *k*th of *K* such random orderings, after computing Spearman’s rank correlation *ϕ*^(*k*)^, we obtain the corresponding Pearson correlation *r*^(*k*)^ using 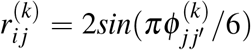. The correlation matrix 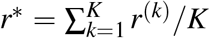 is corrected to be positive definite by setting negative eigenvalues to a small positive value and then renormalizing to preserve the trace of the smoothed correlation matrix. The default choice for MIDASim is *K* = 100. We then take the corrected correlation matrix as the final correlation matrix for the underlying Gaussian model.

To simulate a new dataset with *n* observations, we first generate *n* independent multivariate normal variables 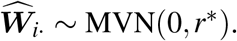 if 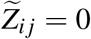 we always choose 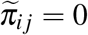.Otherwise, we then choose simulated relative abundances for the *j*-th taxon sampling from the empirical distribution of the non-zero values of *π*_·*j*_. To mimic permutation, if the number of values 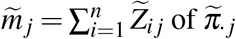 is less than or equal to 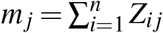, the observed number of zeroes, we sample without replacement; if 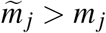 we sample the additional values with replacement, then assign the sampled values so that they agree with the ranking of those *w*_·*j*_ values corresponding to 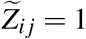.

A count table 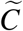 is then calculated by multiplying the sampled relative abundances 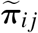 by library size *N*_*i*_ for each observation. Any values so obtained that are between 0 and 1 are rounded up to 1 to keep the presence-absence structure; other values are rounded to the nearest integer. The library sizes for the simulated data are then calculated as 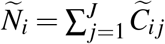 and the final relative abundance is updated through 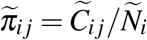.

### 4.3 Parametric Mode using a three-parameter location-scale model for relative abundances

In parametric mode, MIDASim fits the generalized gamma model, a three-parameter distribution in the location-scale family that was proposed for analyzing right-censored survival data [33, 34] to the relative abundance data of each taxon separately. To accomplish this, we define “survival time”

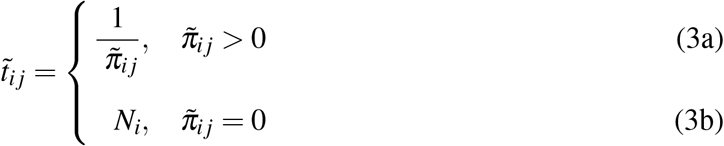

which corresponds to treating 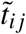 as right-censored when 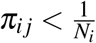. The generalized gamma model then assumes 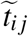 has the distribution specified by

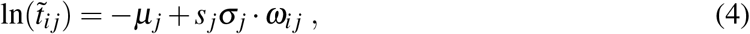

where 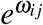 follows a gamma distribution with shape parameter *k*_*j*_ =1*/*|*Q*_*j*_| and scale parameter 1 and where and *s* _*j*_ = sign(*Q*_*j*_). The negative sign on *µ*_*j*_ in (4) is chosen to ensure that the sign of *µ*_*j*_ is positive in a log-linear model for 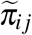. This log-linear model is derived by using Equation (3) in Equation (4).

The resulting cumulative distribution function of 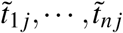 is

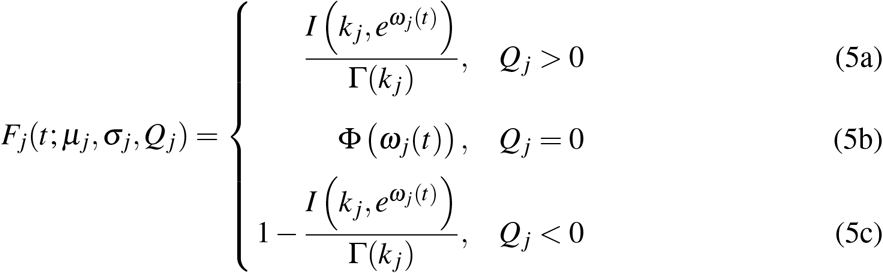

where 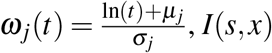 is the lower incomplete gamma function, 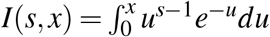, and Γ(·) is the gamma function. Note that log-normal distribution is a special case of the generalized gamma distribution with the scale parameter *Q* = 0.

Although the likelihood for data 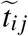 easily accounts for censoring, we found that the maximum likelihood estimators [35] of parameters (*µ*_*j*_, *σ*_*j*_, *Q*_*j*_) gave a poor fit to microbiome data, presumably because for many taxa there are very few non-zero relative abundances. Instead, we developed a novel variant on the method-of-moments approach to estimating these parameters. The *r*^th^ non-central moment of the generalized gamma (for both positive and negative values of *r*) are given [36] by

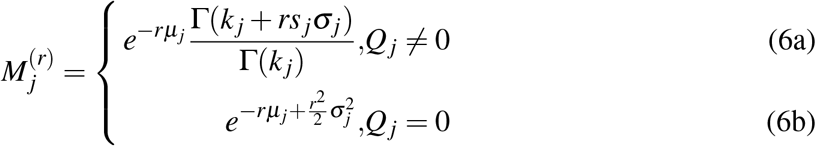

The (empirical) moments of 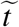 are difficult to estimate because of censoring (i.e., cells having zero counts). However, the empirical moments of 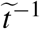 (i.e., the empirical moments of 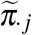) are easily calculated from the template data. For fixed *Q*_*j*_, we can easily find values of 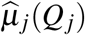 and 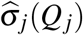 so that the empirical moments of 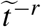 match the theoretical values in (6) for *r* = −1, −2. This task is simplified by the observation that the coefficient of variation (variance/mean^2^) is independent of *µ*_*j*_ which allows determination of 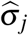 without knowledge of 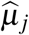 (when *Q*_*j*_ *>* 0 we impose the condition that *σ*_*j*_ *< k*_*j*_*/*2 to ensure the needed moments exist, but can show such a solution always exists). Note these empirical moments are calculated using all observations, not just those having non-zero relative abundance, which stabilizes our approach. To find *Q*_*j*_, we match the observed and expected proportion of zero taxa by maximizing the (profile) likelihood that a zero cell is observed, i.e. we maximize

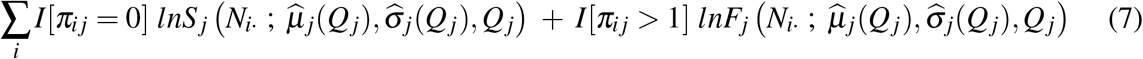

with respect to *Q*_*j*_, where *S* _*j*_(*t*; *µ, σ, Q*) = 1 − *F*_*j*_(*t*; *µ, σ, Q*) is the survival function for the generalized gamma distribution given in (5). Comparison of the predicted and empirical estimates of the CDF of relative abundance for taxa having a wide range of relative abundances are given in Figure S4 and Figure S5).

Given the parameter estimates 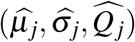, we then generate 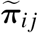 for observations having 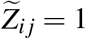 by sampling 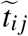 from the generalized gamma distribution upper-truncated at library size *N*_*i*_, then invert 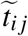 and normalize to obtain 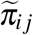 as specified in (3).

The (marginal) predicted probability of being non-zero of *i*-th subject and *j*-th taxon is

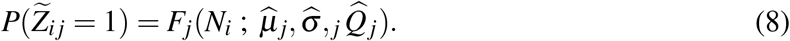

Thus, the predicted number of non-zero cells from *j*-th taxon is 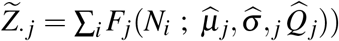. In Figure S6, we show that the empirical (*Z*_·*j*_) and predicted 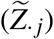 number of non-zero cells are in close agreement. Since the (marginal) probability of being non-zero is specified by (8), we can sample values 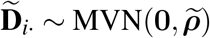 and take 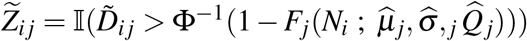, so that (8) is satisfied. Note that estimating *θ*_*j*_ and *η*_*i*_, described in Section 4.1 and used in nonparametric mode, is unnecessary.

### 4.4 Changing the parameters of the simulation

Simulated microbiome data are typically required for rigorous evaluation of methods for analyzing microbiome data. To this end, it is necessary to be able to generate microbiome data sets that are systematically different from the template dataset in a controlled way. In nonparametric mode, users are able to generate data having a different number of samples, different library sizes, different taxon mean relative abundances ***p*** and/or different proportions of zero cells ***δ*** for each taxon. When these changes are made, MIDASim will adjust its marginal distribution quantities and then generate new data having the same presence-absence correlation *ρ* and relative abundance correlation *r*^∗^ as the original data. Note that changes in the mean relative abundance *p*_*j*_ without precisely balanced changes in the taxon proportion of non-zeros *δ*_*j*_ implies changes in the distribution of relative abundances in non-zero taxa, which is used to sample relative abundances for non-zero taxa. In nonparametric mode, MIDASim calculates the mean relative abundance of non-zero cells as 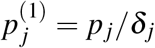, then finds the value *α*_*j*_ for each taxon such that 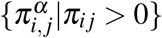 has mean 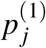 for each taxon. Further, because the number of zero cells in a sample is related to its library size, in nonparametric mode, if users wish to change library sizes, they must also specify the values of *m*_*j*_ and *n*_*i*_ for use in (1) and (2).

Unfortunately, the freedom given in the nonparametric mode may be difficult to use in a controlled simulation study. For example, if we wish to change the library sizes of certain observations or the relative abundances of various taxa, it is not clear how the proportion of non-zero taxa should change. This is where the parametric mode of MIDASim is most useful, as changes in the parameters of the parametric model (including library sizes) imply coordinated changes in all other quantities. For example, the proportion of non-zero cells for each taxon is given by (8), which facilitates changing library sizes if desired. Because the model used for relative abundance in parametric mode is a log-linear model in the location-scale family, changes in taxon relative abundance can achieved directly by changing the parameters *µ*_*j*_ while holding other parameters fixed. Note that *µ*_*j*_ is the mean on the log scale; the mean on the relative abundance scale is given by (6). For convenience, MIDASim in parametric mode allows the user to specify a new value of the taxon mean relative abundances *p*_*j*_ and will convert these values to the corresponding values of *µ*_*j*_ assuming 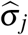 and 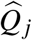 are unchanged.

After either modification of the parameters, we predict the number of non-zero cells in each subject 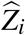 and that in each taxon 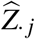 using (8), and then use the marginal totals 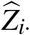 and 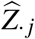 in (2) and (1) for use in generating the presence-absence data 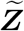. In either mode, once 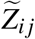 is obtained, changing the number of samples is easily accomplished by simply generating extra observations using the copula model.

In summary, MIDASim takes an OTU count table as input, and output simulated tables of counts, relative abundances and presence-absence data. Its nonparametric mode permits adjustments in sample size, library sizes, mean relative abundances, and the proportion of non-zero cells. These alterations in the nonparametric mode affect simulations in two ways: firstly, changes to sample size, library sizes, and the proportion of non-zero cells directly influence the values of *m*_*j*_ and *n*_*i*_ in Equations (1) and (2), thereby altering the construction of the presence-absence matrix; secondly, variations in mean relative abundances lead to recalibrations in the values of non-zero relative abundances, impacting the empirical marginal distribution of these abundances. In contrast, the parametric mode offers coordinated changes, allowing for adjustments in library sizes, mean relative abundances, and the location parameters ***µ*** in the generalized gamma model. Alterations in mean relative abundances are reflected in the estimation of ***µ*** to align with the first moment, leading to distinct generalized gamma models. Similarly, adjustments in library sizes affect the predicted probability of a non-zero presence, as determined by Equation 8, which influences both *m*_*j*_ and *n*_*i*_ values and consequently the structure of the presence-absence matrix.

### 4.5 Assessment of Differential Abundance Analysis Methods using MIDASim-Simulated Data

We used MIDASim in parametric mode to simulate *n* = 100 and *n* = 200 independent microbiome samples using the IBD data as the template. For each observation we simulated two binary covariates *X*_1_ and *X*_2_ in such a way that the covariates divide the sample into four equal-sized groups. The group having *X*_1_ = *X*_2_ = 0 was the “null” or control group. To model the effect of covariates in the other groups, we randomly selected either *M*_1_ = 10 or *M*_1_ = 20 “causal” taxa from the top 100 most abundant taxa to exhibit differential abundance based on *X*_1_. Additionally, we selected a set of *M*_2_ = 10 “causal” taxa showing differential abundance based on *X*_2_, with an overlap of 5 taxa between the two sets of causal taxa. Fitting MIDASim to the template data provided 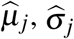 and 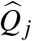 for each taxon. For the non-null groups, we modified the values of *µ*_*j*_ according to the model

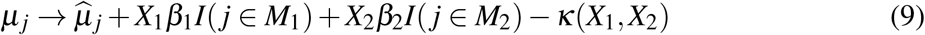

where *κ*(*X*_1_, *X*_2_) is chosen so that the resulting mean relative abundances are normalized for each choice of covariates. This corresponds to choosing mean relative abundances in the non-null groups to be

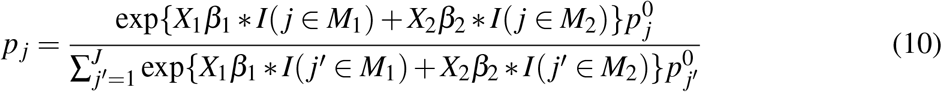

where 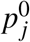 is the mean relative abundance for taxon *j* in the null (template) data.

We varied *β*_1_ from 0.5, 1, 1.5, 2, and *β*_2_ was fixed at 1 (corresponding to treating *X*_2_ as a confounder whose effect size is not of interest). We used MIDASim to generate data from each covariate group, using the same values of *ρ* (tetrachoric correlation matrix) and *r*^∗^ (copula correlation matrix) as in the null (template) data. Library sizes for each covariate group were sampled with replacement from the set of library sizes in the template data. Relative abundances were calculated using the modified values of *µ*_*j*_ given in (9). False discovery rates (FDR) are based on 500 simulated datasets, based on a nominal value of FDR=0.2.

## Supplementary Information

### Supplementary Files

Currently found at the end of this document

### Authors’ contributions

MH contributed to the development of the method, performed simulation studies and comparisons, and wrote the manuscript. GAS conceived the study, primarily developed the method, and wrote the manuscript. NZ conceived the study, contributed to the development of the method, and wrote the manuscript. All authors read and approved the final manuscript.

## Availability of data and materials

The R package MIDASim is available on GitHub at https://github.com/mengyu-he/MIDASim.

All template datasets are publicly available and can be accessed through R package `HMP2Data`. Details can be found in the vignette of R package `MIDASim`.

## Ethics approval and consent to participate

Not applicable.

## Consent for publication

Not applicable.

## Acknowledgements

Not applicable.

## Author affiliations

Department of Biostatistics and Bioinformatics, Emory University, Atlanta, 30322, GA, USA (Mengyu He)

Department of Gynecology and Obstetrics, Emory University, Atlanta, 30322, GA, USA (Glen A. Satten)

Department of Biostatistics, Johns Hopkins University, Baltimore, MD 21205, USA (Ni Zhao)

## Supplementary File: Statistical Analyses

We compared the simulated data from each method to the template data using several measures. First, we concatenated the template data with a simulated dataset from each method, and defined a binary variable to differentiate the template and simulated data. We tested the significance of this variable using PERMANOVA [24], which tests for shifts in the between-observation distances. Our PERMANOVA tests used the Jaccard distance as well as the Bray-Curtis distance, which are both commonly used in microbiome data analyses. The Jaccard distance uses only presence-absence information in the data, and thus can assess how similar 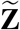 and **Z** are, while the Bray-Curtis distance accounts for both the presence-absence and relative abundance information and can be used to assess the simulation of 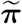. We also compared the alpha diversity of the simulated data and template data. The simulated communities were compared to the template in terms of observed richness and Shannon Index, and the differences in diversity were tested by Kruskal-Wallis tests. The observed richness is simply the number of observed taxa, while Shannon Index additionally considers evenness-the relative abundances of taxa-when quantifying diversity. To suppress random variability, we repeated the comparison of alpha-diversity and beta-diversity using 20 simulated datasets from each of the four methods. Finally, we compared the methods visually, using ordination and PCoA, as well as boxplots of alpha diversity values, using a single simulated data set for each method.

We next compared the simulation approaches in terms of their *β* -dispersion, by comparing whether the distribution of distances from each observation to the sample centroid was the same in the simulated and template data. We calculated distances to the centroids using the betadisper function in R package vegan [37]. We used the Kolmogorov-Smirnov (K-S) test to compare these empirical distributions. We again averaged results over 20 simulation replicates to suppress random variability. We also compared the alpha diversity of the template and simulated data, as measured by the species richness (number of observed taxa) and the Shannon entropy.

Finally, we evaluated the performance of our approach to generating data with different library sizes by rarefying our template datasets, then using the approach described in section 2.3 to increase the library size to that of the original template data. Thus, we can compare the resulting simulated data to the original template data. Specifically, for each template, the observed counts for each subject were rarefied (subsampled without replacement) to remove 10% of the observed counts. The rarefied data are then treated as the template data in MIDASim, and the target library size is the original library size.

## Supplementary File: Tables and Figures

**Table S1:**
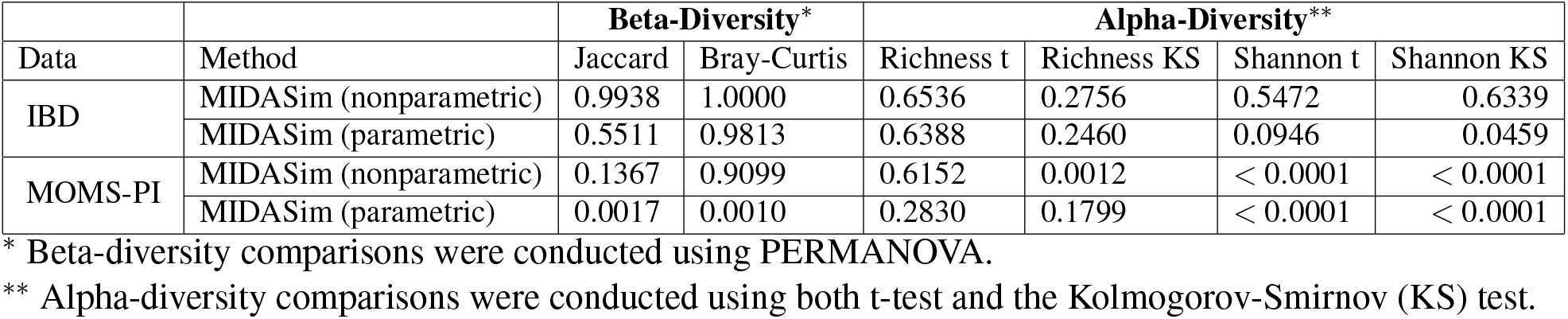
Average *p*-values for comparing alpha and beta diversities in MIDASim simulated data (20 replicates) versus template data, without removal of rare taxa.

**Table S2:** Summary statistics of the IBD and MOMS-PI datasets used in comparison after filtering.

| Dataset | Sample size | # of taxa | Log10 Library size<br>mean (min, max) | % of zeros | CV*<br>mean (min, max) |
| --- | --- | --- | --- | --- | --- |
| IBD | 146 | 614 | 4.22 (3.51, 4.50) | 85.09 | 6.24 (0.90, 11.98) |
| MOMS-PI | 517 | 1146 | 4.61 (3.50, 5.78) | 95.25 | 13.58 (1.65, 22.72) |

\* CV is the coefficient of variation of observed OTU counts for each taxon.

**Table S3:** Summary of CPU time and memory usage for fitting templates and simulating one dataset with varying taxa (*J*) and sample size (*n*). Template sizes range from 100 to 1000 taxa, and sample sizes vary between 100 and 5000. Simulated datasets match the size of the corresponding templates in each *J* and *n* combination.

| Mode | Sample size | Time $s$ | | | Memory allocation (MB) | | |
| --- | --- | --- | --- | --- | --- | --- | --- |
| | | $J = 100$ | $J = 500$ | $J = 1000$ | $J = 100$ | $J = 500$ | $J = 1000$ |
| nonparametric | $n = 100$ | 3.3 | 18.8 | 57.8 | 182.4 | 1261.2 | 3212.0 |
| | $n = 1000$ | 16.4 | 105.7 | 337.5 | 1529.2 | 6781.2 | 16574.8 |
| | $n = 5000$ | 73.3 | 517.6 | 1606.0 | 8138.9 | 40427.4 | 82572.2 |
| parametric | $n = 100$ | 4.3 | 25.2 | 70.1 | 190.1 | 1298.4 | 3262.3 |
| | $n = 1000$ | 15.4 | 111.4 | 338.8 | 1509.7 | 8220.8 | 17454.1 |
| | $n = 5000$ | 71.4 | 526.0 | 1569.5 | 7768.5 | 39969.2 | 81411.3 |

**Figure S1:**
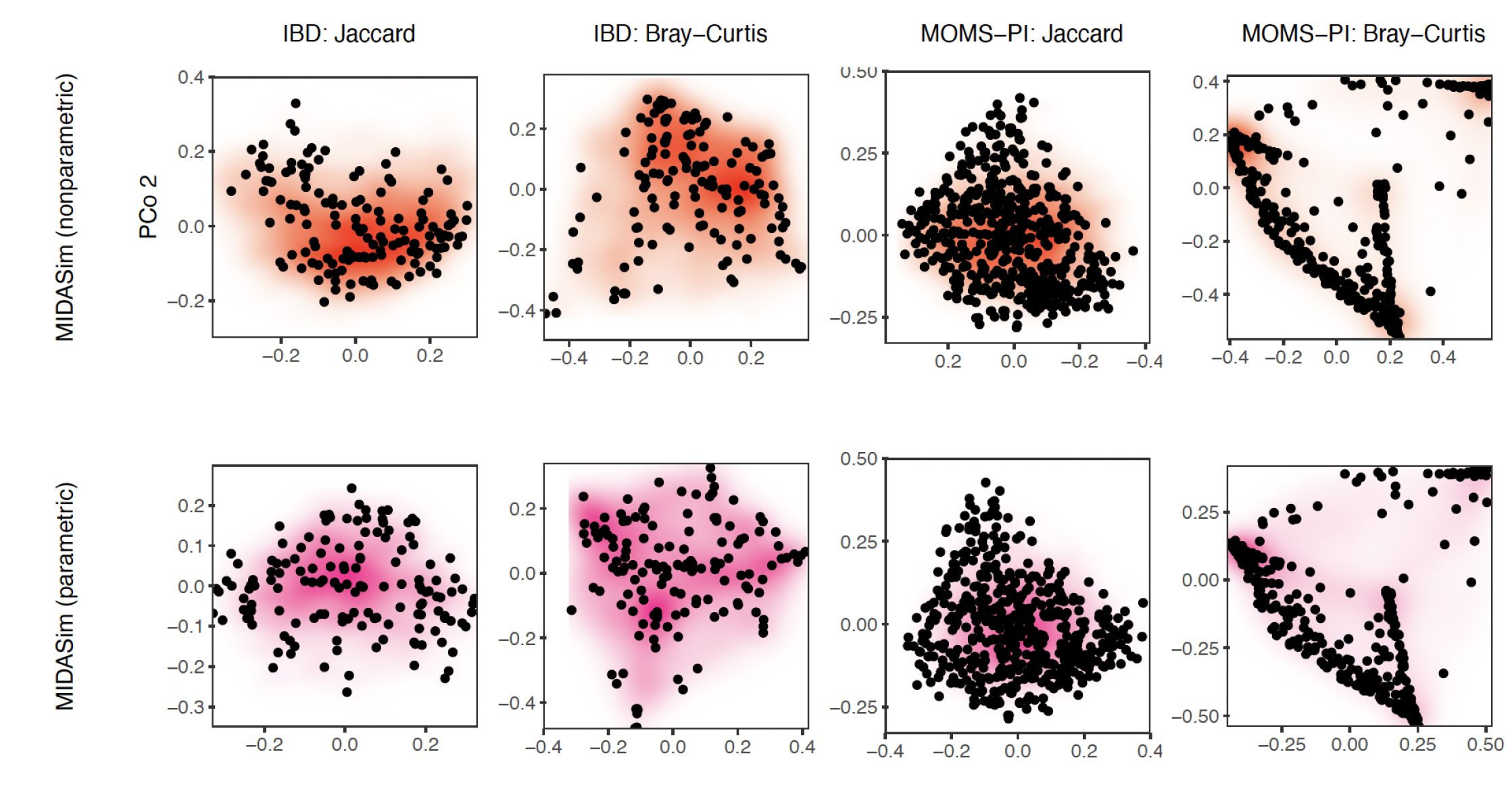
Principal Coordinates plots (PCoA) of the simulated and original microbiome community. The colored density map is plotted based on 20 replicates of simulated communities by MIDASim, with darker coloring associated with higher density of simulated values. Black points represent the original community.

**Figure S2:**
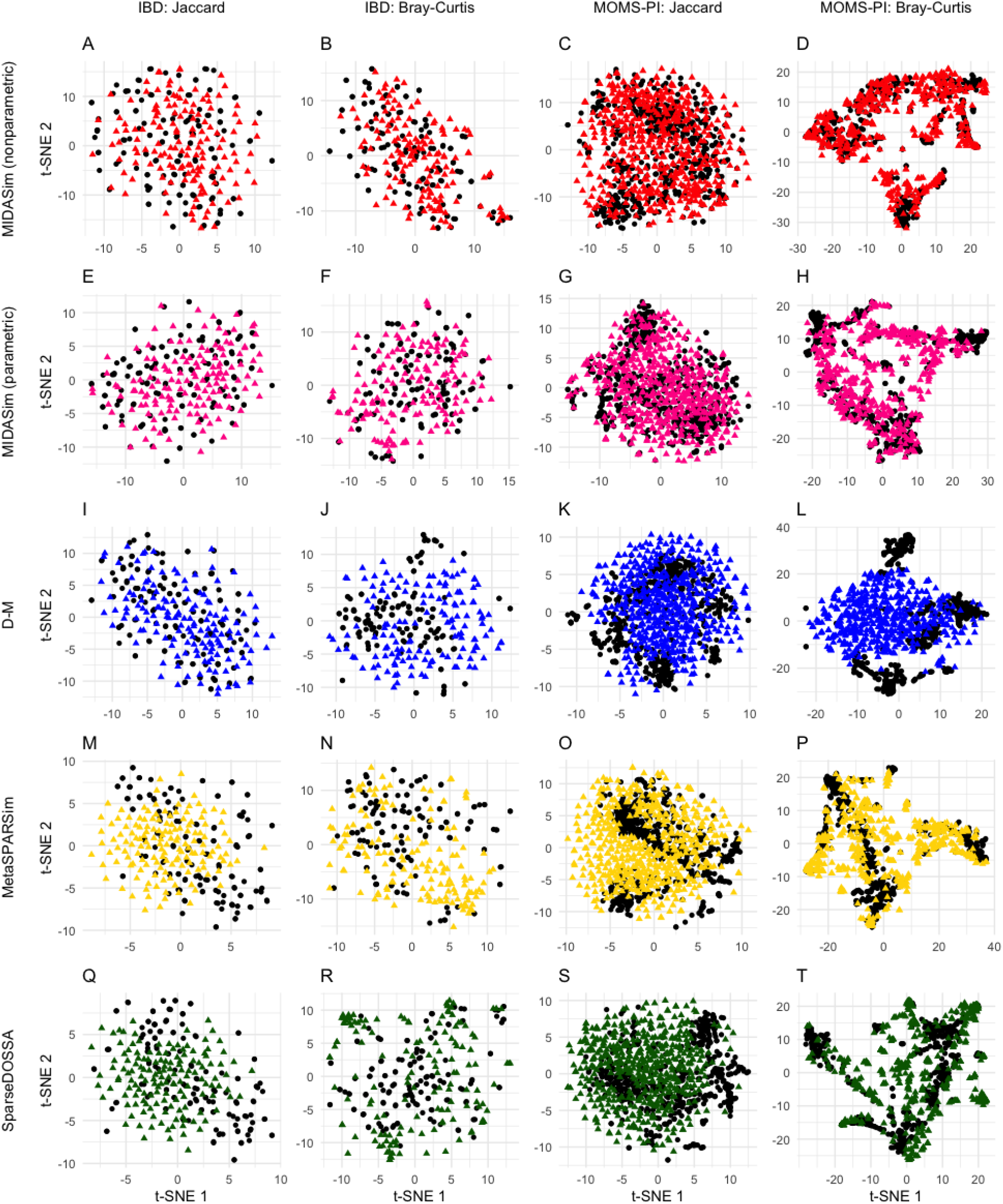
Plots of t-distributed stochastic neighbor embedding (t-SNE) of the simulated and original community. Each row corresponds to one method. The left two columns are the plots for the IBD data, and the right two columns are the plots for the MOMS-PI data. Black points: samples from original data. Colored points: samples from the simulated data with red being MIDASim with nonparametric model, pink being MIDASim with parametric model, blue being D-M, yellow being MetaSPARSim, and green being SparseDOSSA.

**Figure S3:**
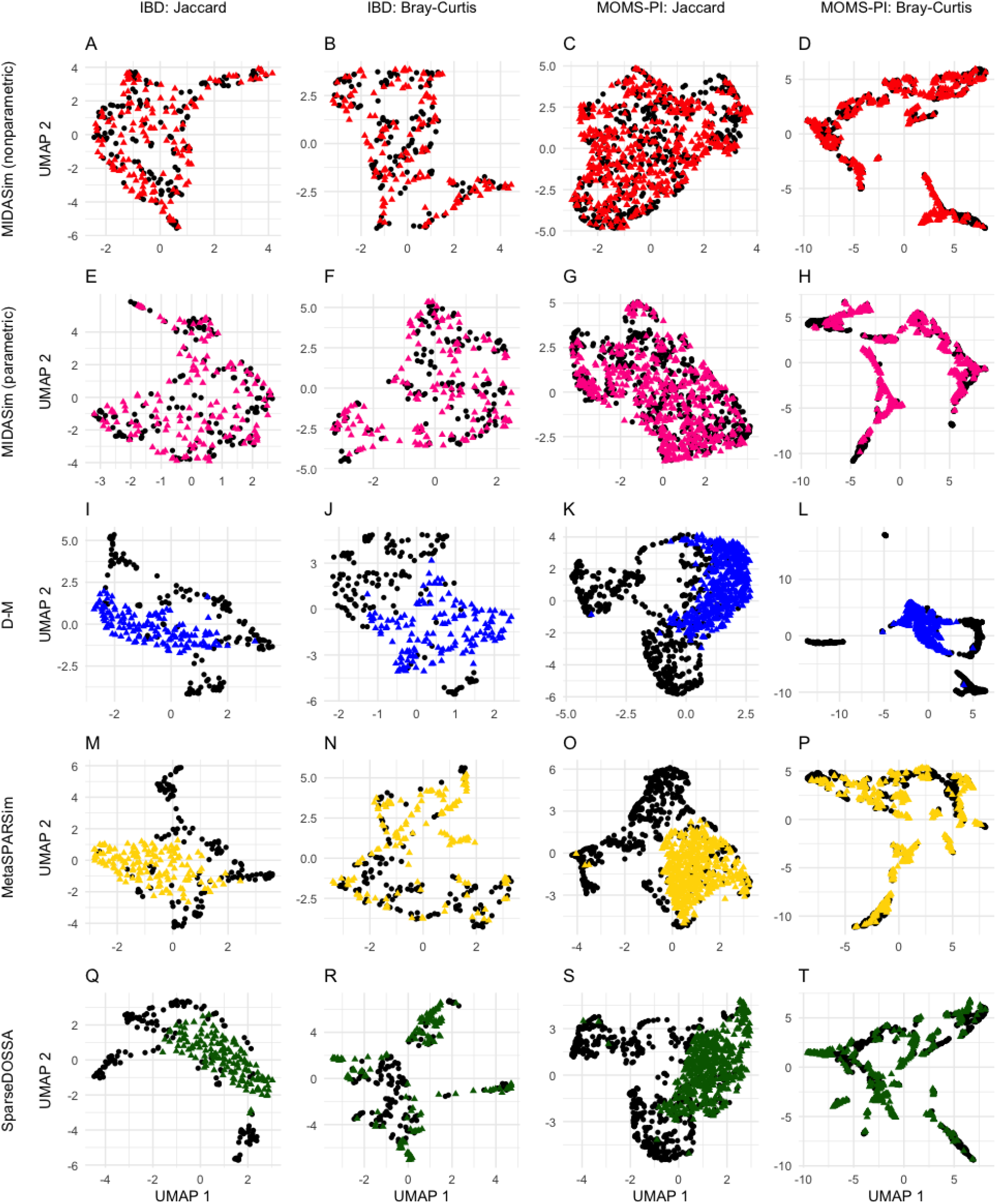
Plots of Uniform Manifold Approximation and Projection (UMAP) of the simulated and original community. Each row corresponds to one method. The left two columns are the plots for the IBD data, and the right two columns are the plots for the MOMS-PI data. Black points: samples from original data. Colored points: samples from the simulated data with red being MIDASim with nonparametric model, pink being MIDASim with parametric model, blue being D-M, yellow being MetaSPARSim, and green being SparseDOSSA.

**Figure S4:**
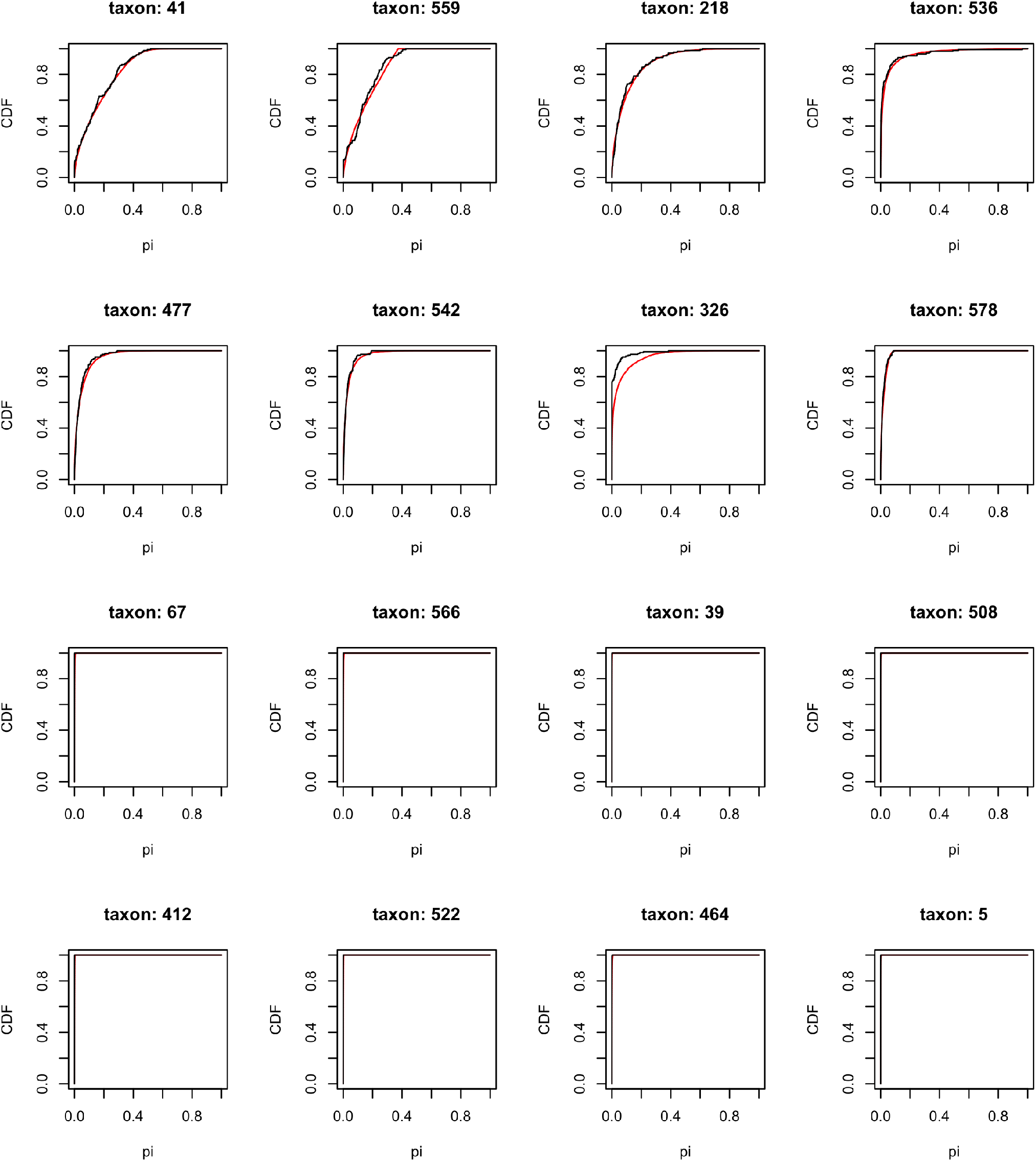
Comparison of the predicted (red) and empirical (black) estimates of the CDF of relative abundance for the top 8 and moderately abundant 8 taxa in IBD dataset.

**Figure S5:**
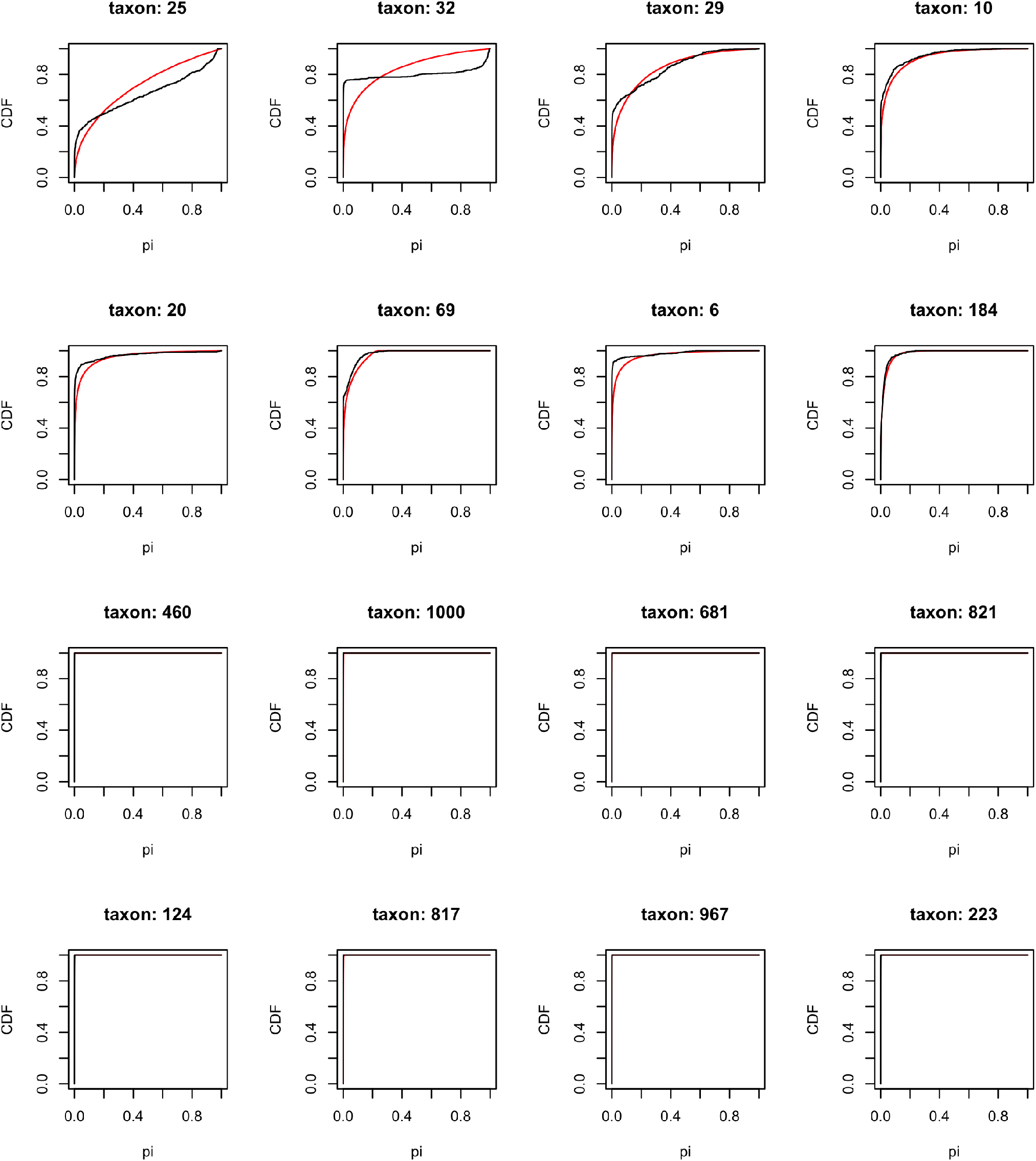
Comparison of the predicted (red) and empirical (black) estimates of the CDF of relative abundance for the top 8 and moderately abundant 8 taxa in MOMS-PI dataset.

**Figure S6:**
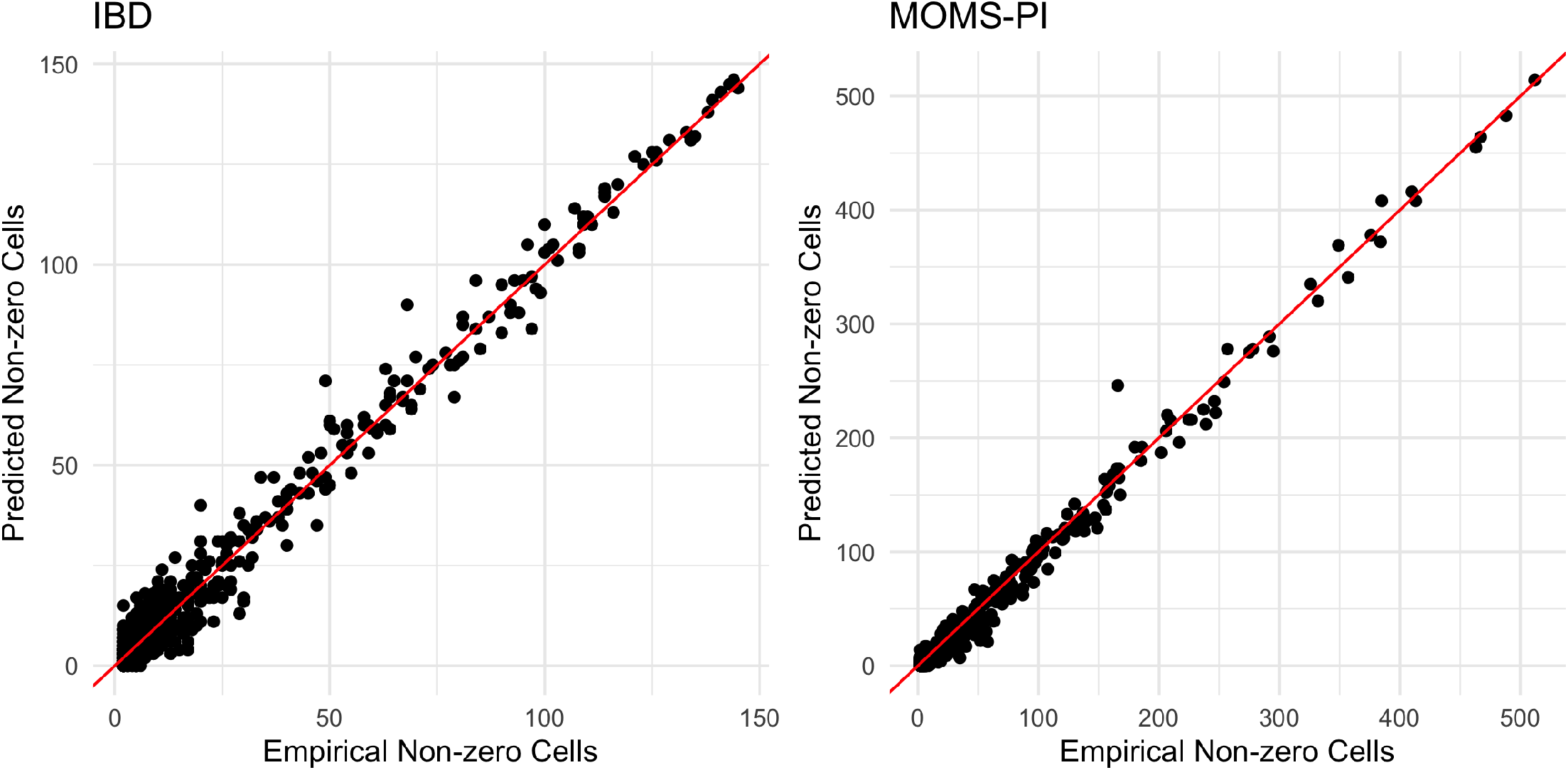
Comparison of the empirical *Z*_*j*_ and predicted 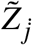 number of non-zero cells in IBD and MOMS-PI datasets. The red lines represent the diagonal reference lines.

